# Knockout of *Hsp70* genes significantly affects locomotion speed and gene expression in leg skeletal muscles of *Drosophila melanogaster*

**DOI:** 10.1101/2023.12.01.569551

**Authors:** Pavel A. Makhnovskii, Inna V. Kukushkina, Nadia S. Kurochkina, Daniil V. Popov

**Affiliations:** Institute of Biomedical Problems of the Russian Academy of Sciences, Moscow, 123007, Russia; Lomonosov Moscow State University, Moscow, 119234, Russia

**Keywords:** heat shock proteins, skeletal muscle, transcriptome, exercise, training

## Abstract

The functions of the *Hsp70* genes were studied using a line of *D. melanogaster* with knockout of six these genes out of thirteen. Namely, effect of knockout of *Hsp70* genes on negative geotaxis climbing (locomotor) speed and the ability to adapt to climbing training (0.5-1.5 h/day, 7 days/week, 19 days) were examined. Seven- and 23-day-old *Hsp70^−^* flies demonstrated a comparable reduction (2-fold) in locomotor speed and widespread changes in leg skeletal muscle transcriptome (RNA-seq), compared to *w^1118^*flies. To identify the functions of genes related to decreased locomotor speed the overlapped differentially expressed genes at both time points were analyzed: the up-regulated genes encoded extracellular proteins, regulators of drug metabolism and antioxidant response, while down-regulated genes encoded regulators of carbohydrate metabolism and transmembrane proteins. Additionally, in *Hsp70^−^* flies, activation of transcription factors related to disruption of the fibril structure and heat shock response (Hsf) were predicted, using the position weight matrix approach. In the control flies, adaptation to chronic exercise training was associated mainly with gene response to a single exercise bout, while the predicted transcription factors were related to stress/immune (Hsf, NF-kB, etc.) and early gene response. In contrast, *Hsp70^−^* flies demonstrated no adaptation to training, as well as significantly impaired gene response to a single exercise bout. In conclusion, the knockout of *Hsp70* genes not only reduced physical performance, but also disrupted adaptation to chronic physical training, which is associated with changes in leg skeletal muscle transcriptome and impaired gene response to a single exercise bout.

**New & Noteworthy:**

- Knockout of six *Hsp70* genes in *D. melanogaster* reduced locomotion (climbing) speed that is associated with genotype-specific differences in leg skeletal muscle gene expression.
- Disrupted adaptation of *Hsp70^−^* flies to chronic exercise training is associated with impaired gene response to a single exercise bout.

## Introduction

Skeletal muscle tissue plays a key role in regulating carbohydrate and fat metabolism as well as insulin sensitivity in the organism. Maintaining oxidative capacity and skeletal muscle mass (strength) is key to maintaining physical performance and quality of life (1–3). Inactivity and disuse lead to a sharp decrease in oxidative capabilities, insulin sensitivity, and physical performance of skeletal muscles, while chronic physical exercise (in particular endurance exercise) quickly restores/increases these parameters. This increase is related to the expression of genes encoding regulators of carbohydrate and fat metabolism, energy (in particular, oxidative) metabolic enzymes, and the transcript-specific translation of mRNAs encoding mitochondrial proteins (reviewed in (4)). More than 99% of mitochondrial proteins are encoded by genomic DNA and synthesized in the cytoplasm as preproteins, therefore the transport of these proteins into mitochondria, their processing and folding must play an important role in the exercise-induced activation of mitochondrial biogenesis (5, 6).

Heat shock protein (Hsp) family 70 (∼10 genes) and 90 (∼4 genes) play an important role in the intracellular transport and import of mitochondrial proteins (6). A number of studies have examined effects of Hsp70 (encoded by the *Hspa* family genes) on skeletal muscle. It was shown that overexpression of *Hspa1a* in skeletal muscles activated mitochondrial biogenesis and increased aerobic (endurance) performance and insulin sensitivity in mice (7, 8). Similarly, overexpression of *Hspa9*, encoding a mitochondrial chaperone, in cultured cardiomyocytes and mouse hearts increases the content and activity of oxidative enzymes and mitochondrial function, as well as deactivates the production of reactive oxygen species and the apoptotic program (9, 10). These findings are consistent with data showing an increase in the content of various Hsp and mitochondrial proteins after chronic aerobic training (11–13). Furthermore, increased expression of the *Hspa1a* gene accelerates muscle regeneration after damage (reviewed in (14)) and prevents a decrease in muscle mass and strength induced by disuse (15–17) and dystrophy (18), indicating the pleiotropic effects of *Hsp70*. The *Hspa*-dependent effects may be related to the increase in the phosphorylation/activation of mitogen- and AMP-activated kinases (JNK and AMPK), expression of deacetylase SIRT1 and mitochondrial transcription factor *Tfam*, as well as deactivation of transcription factors NF-kB and FOXO3a and genes encoding muscle-specific E3 ligases *Trim63* and *Fbxo32* (7, 8, 15, 17). It should be noted that the precise mechanism(s) underlying the *Hsp*-dependent effects (in particular, regulation of gene expression) are still elusive.

Loss-of-function experiments play an important role in understanding gene function. Suppression of one of the *Hsp70* (*Hpsa*) genes can be compensated by other genes from this family, which makes it difficult to interpret the results of such experiments. In the last decade, flies have been extensively used as a model for studying the mechanisms of adaptation to physical exercises (reviewed in (19)). In the present work, the functions of the *Hpsa* (*Hsp70*) genes were studied using a line of *D. melanogaster* with knockout (deletion) of six *Hsp70* genes (orthologs of the mammalian genes *Hspa1a*, *Hspa1b*, *Hspa2*, and *Hspa8*) (20) out of thirteen. Namely, the authors assessed effect of knockout of *Hsp70* genes on negative geotaxis climbing (locomotor) speed, a marker of endurance performance, and on the ability to adapt (i.e., improve climbing speed) to increased physical activity (climbing training 0.5-1.5 h/day, 7 days/week, 3 weeks). The *Hsp70*-dependent changes in transcriptomic profiles in fly legs, consisting mainly of muscle bundles (21), were examined after chronic exercise (climbing) training and a single exercise bout; then transcription factors associated with changes in gene expression were predicted using the position weight matrix (PWM) approach.

## Methods

### D. melanogaster strains and cultivation conditions

Males *D. melanogaster* adapt to chronic exercises better than females (22). Therefore, *w^1118^*males (Vienna Drosophila Resource Center, ID 60000) and *Hsp70^−^*males (genotype *w^1118^*; *Df(3R)Hsp70A*, *Df(3R)Hsp70B*, Bloomington Drosophila Stock Center, ID 8841) with a knockout of 6 genes (*Hsp70Aa*, *Hsp70Ab*, *Hsp70Ba*, *Hsp70Bb*, *Hsp70Bbb*, *Hsp70Bc*) (20) were used in this study. *Hsp70^−^* flies have two deletions located in 87A (*Hsp70Aa* and *Hsp70Ab*) and 87C (*Hsp70Ba*, *Hsp70Bbb*, *Hsp70Bb*, and *Hsp70Bc*) loci of chromosome 3R. Deletions were obtained by homologous recombination with the flanking regions using the linearized (by I-SceI endonuclease) P-element vector (pW25) and FLP recombinase. Hypomorphic *w^hs^* gene was used as marker of deletion, subsequent deletion of *w^hs^* was obtained by Cre-mediated recombination. The lack of expression of these genes was verified by analysis of their open reading frame coverage (RNA sequencing) in a previous study (23). Flies were kept at 22 °C, humidity 45–55%, with a 12-hour light-dark cycle, on a food medium containing 5% sucrose, 10% baker’s yeast, 5% semolina, 0.7% agar, and 0.1% propionic acid. For all experiments, 30 males were cultivated in 50-ml polypropylene tubes with a vented lid, which contained 5 ml of food medium.

After eclosion, males of each strain were divided into 4 groups: 2 groups for an experiment with chronic training (trained and untrained groups) and 2 groups for an experiment with a single exercise bout (exercised and unexercised groups) (Figure 1).

**Figure 1.**
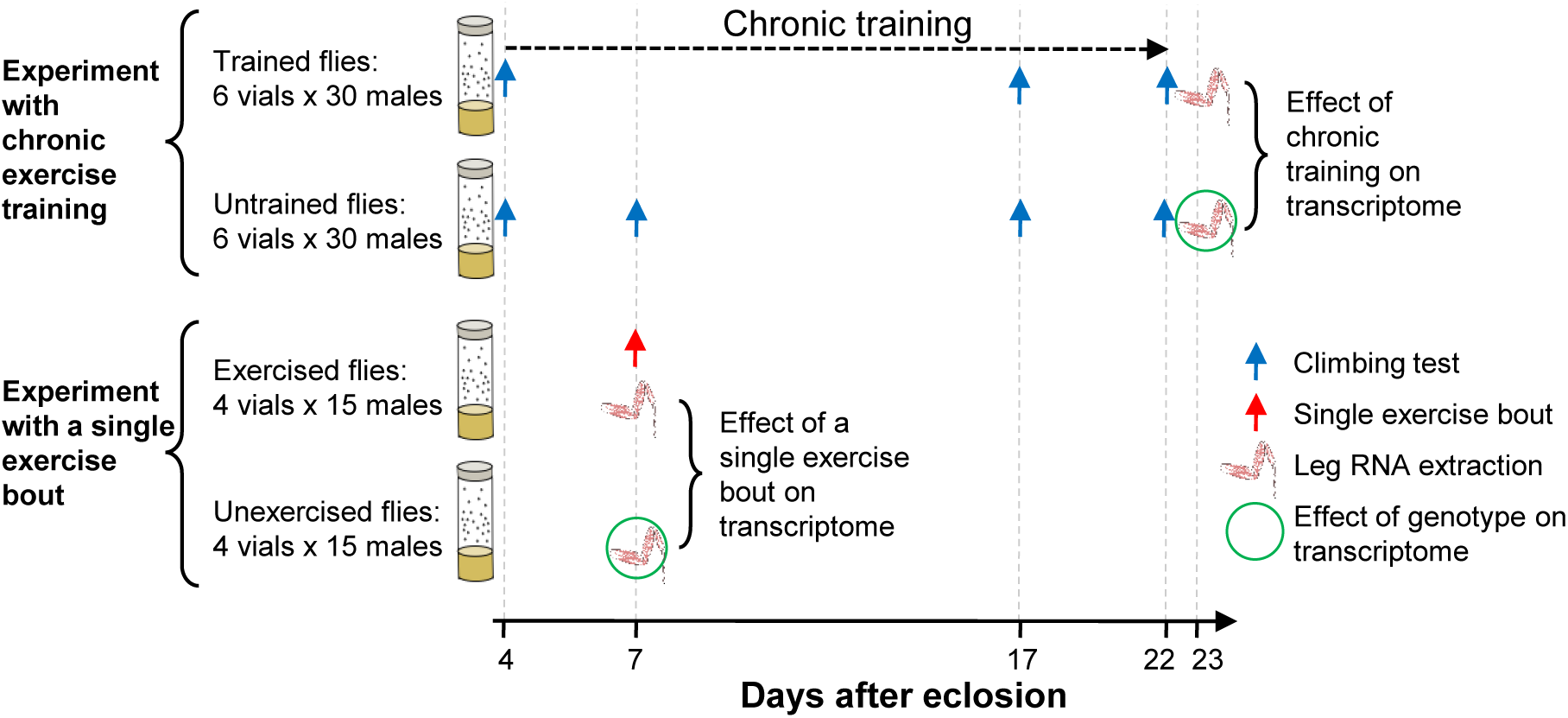
Design of the study (for each strain).

### Climbing (locomotion) speed

Negative geotaxis climbing speed was evaluated using a homemade device that rotated the tubes 180 degrees every 15 s around the transverse axis as previously described (24). Five minutes after the start of the test, a video was recorded (for 6 min) to determine the position and climbing speed of each fly using an automated measurement software (FreeClimber tool (25)).

### Chronic exercise (climbing) training

Flies (n = 180 for each strain) were trained from the 4th to the 22nd day after eclosion (19 days) using a homemade device, which rotated the tubes every 15 s as described above. Each training session began at 12:00; the training duration was 30 min on the first 2 days, 60 min on the second 2 days, and 90 min on the remaining days. Unexercised flies (n = 180 for each strain) were kept in 1.5 cm height tubes as previously described (24). Flies were anesthetized with diethyl ether 24 h after the last training session; legs were removed in phosphate-buffered saline and then placed in RNA extraction buffer (see below).

### Single bout of exercise (climbing)

Untrained flies (7 days old, n = 60 for each strain) exercised for 2 h as described above. Immediately thereafter, exercised and unexercised flies were anesthetized and legs were removed for RNA extraction as described above.

### RNA Sequencing and Data Processing

The protocol was described previously (23). Namely, legs from 15 flies were homogenized in the chilled RLT buffer (Qiagen, Germany) with 1% beta-mercaptoethanol in a 1.5-ml tube using a polypropylene pestle and a drill and then incubated (55 °C, 10 min) with protein kinase K (Evrogen, Russia). Total RNA was extracted by a silica spin column (CleanRNA Standard, Evrogen, Russia); RNA concentration was evaluated by a fluorimetric assay (Qubit 4, Thermo Scientific). RNA integrity was evaluated in an aliquot of total RNA (200 μg) after DNase I (Thermo Scientific) treatment using capillary electrophoresis (TapeStation, Agilent, Germany). Five hundred micrograms of total RNA was used to prepare strand-specific libraries by the NEBNext Ultra II Directional RNA Library Preparation kit (NEB) and sequenced (75 nucleotides, single end) with a median depth of 25 million reads per sample (NextSeq 550, Illumina, USA).

Data processing was performed, as described previously (23). Low-quality reads and adapter sequences were deleted (Timmomatic tool, v 0.36), and then the reads were aligned to the BDGP6.94 primary genome assembly. Uniquely aligned reads were counted for known exons of each gene using the Rsubread package (R environment) and Ensembl annotation (BDGP6.94). Then low expressed genes with TPM <1 were removed (kallisto v0.46.2) and differential expression analysis for protein-coding genes was performed by the DESeq2 method (analysis of unpaired samples with the Benjamini-Hochberg correction). Differentially expressed genes were defined as protein-coding genes with |fold change| >1.25 and p_adj_ <0.05.

### Transcription factor prediction

To search for transcription factors associated with transcriptomic changes, the PWM approach was used, as described previously (26). Namely, enrichment of predicted transcription factor binding sites (and corresponding transcription factors) in promoters (–300 to +100 b.p. from the transcription start site of the most abundant mRNA isoform) was performed by the GeneXplain platform (the “Search for enriched TFBSs (tracks)” function http://wiki.biouml.org/index.php/Search_for_enriched_TFBSs_(tracks)_(analysis)) using the PWM database TRANSFAC v2020.3 (“insects”). The maximum enrichment (FE_adj_, statistically corrected odds ratios with a confidence interval of 99%) was determined for each PWM (site frequency ≤1 per 2,000 bp) relative to that in 5,000 random individual promoters showing no differential expression in any of the experimental points (DESeq2 method, p_adj_ >0.4). Adjusted fold enrichment (FE_adj_) >1.0 for transcription factor binding site or promoter’s sequence number (the binomial test and exact Fisher’s test, respectively) and FDR <0.05 were set as significance thresholds. If a TF had several PWMs, the most enriched PWM was used.

### Statistical Analysis

Physiological data were expressed as the median and interquartile range. The Mann-Whitney test with a significance level of 0.05 was used to compare variables. To analyze repeated measures data, the three-way (time, genotype, and training) mixed effects model with Tukey’s multiple comparisons test (significance level 0.05) was used. The functional enrichment analysis (GENE ONTOLOGY BP/CC DIRECT and KEGG PATHWAY databases) was performed relative to all expressed (TPM >1) protein-coding genes using the DAVID 6.8 (p_adj_ <0.05 Fisher exact test with Benjamini correction). To group genes into various functional classes, the FlyBase gene database was used. Networks of protein-protein and genetic interactions were constructed with Cytoscape v.3.10.1 and GeneMANIA using the FlyBase interaction databases and Genemania “Costello-Andrews-2009” set for predicted interactions.

## Results

### Knockout of the Hsp70 genes reduced locomotion speed and disrupted adaptation to exercise training

In *w^1118^* flies, climbing speed rapidly drops after three weeks of life (23, 27, 28); therefore, 7- and 23-day-old flies were used to study effect of *Hsp70* gene knockout. Knockout of six *Hsp70* genes significantly (2-fold) reduced climbing speed at both time points (Figure 2). This is consistent with the previous work on this fly strain (23).

**Figure 2.**
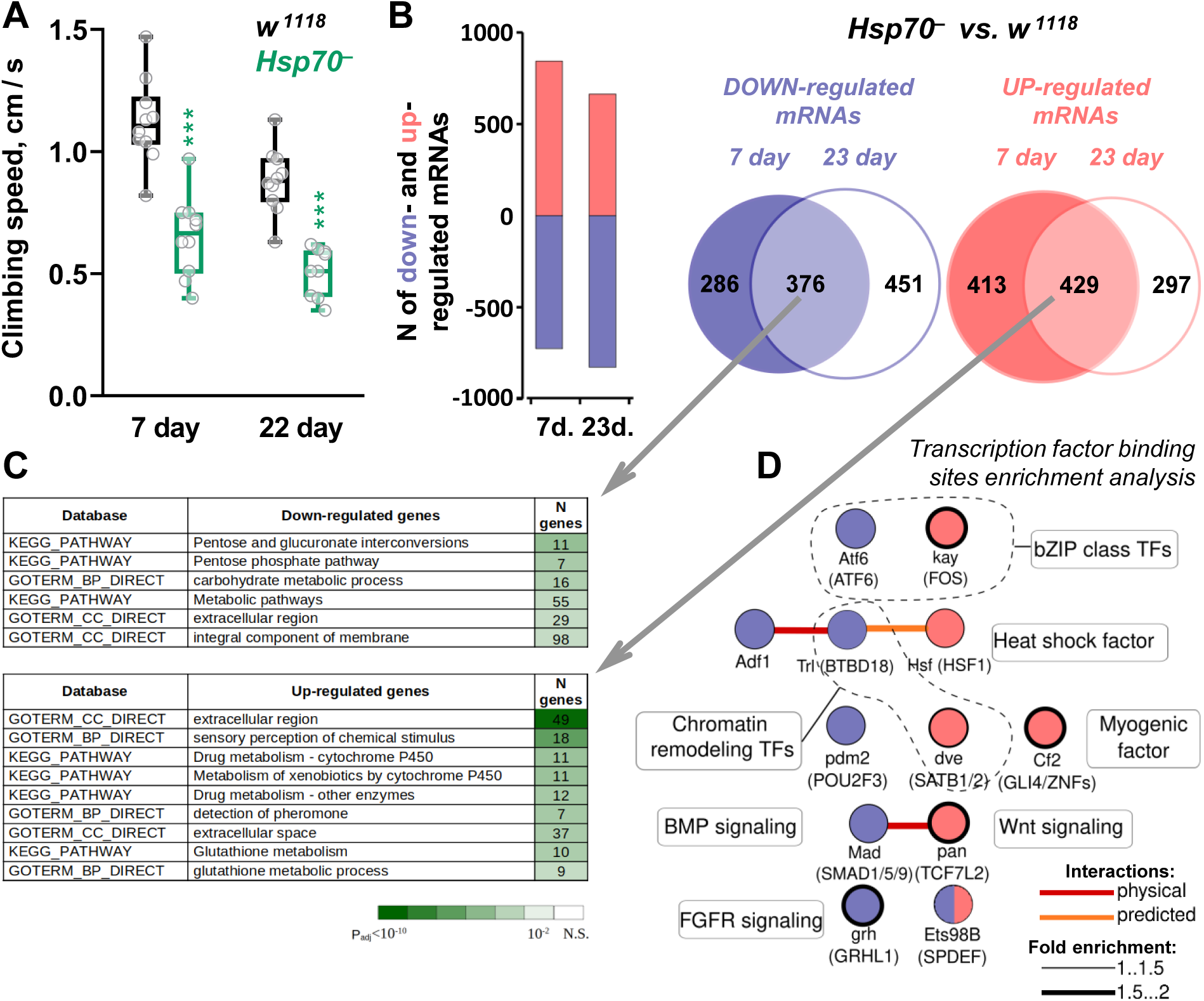
Reduced locomotion (climbing) speed in *Hsp70^−^* flies is associated with genotype-specific differences in leg skeletal muscle gene expression. A – The climbing speed of 7- and 23-day-old *Hsp70^−^* flies was noticeably lower than in the control (*w^1118^*). n = 10 pools (20–30 flies per pool) for each strain. *** – difference from the control at p <0.001. B – Number of up- and down-regulated mRNAs in 7- and 23-day-old *Hsp70^−^* flies compared to the control and their overlap. n = 4 pools (15 flies per pool) for each strain. C – Functions of genes related to reduced locomotion (climbing) speed in *Hsp70^−^*flies, i.e., differentially expressed genes that overlapped between days 7 and 23. The number of genes in each functional category is presented; the heat map shows the p_adj_-value. D – Transcription factors associated with up- and down-regulated genes (red and blue, respectively) that overlapped between days 7 and 23. Lines indicate potential protein-protein interactions; the border thickness indicates the enrichment of the transcription factor binding site in the promoter (–300 to +100 b.p.).

Next, effect of 19-day training on climbing speed was examined. In line with previous works (reviewed in (19)), the training increased the climbing speed in *w^1118^*flies (Figure 3), However, *Hsp70^−^* flies showed no increase in climbing speed induced by chronic training (Figure 3). Together, these findings clearly showed that knockout of the *Hsp70* genes reduced exercise performance and disrupted adaptation to chronic exercise training.

**Figure 3.**
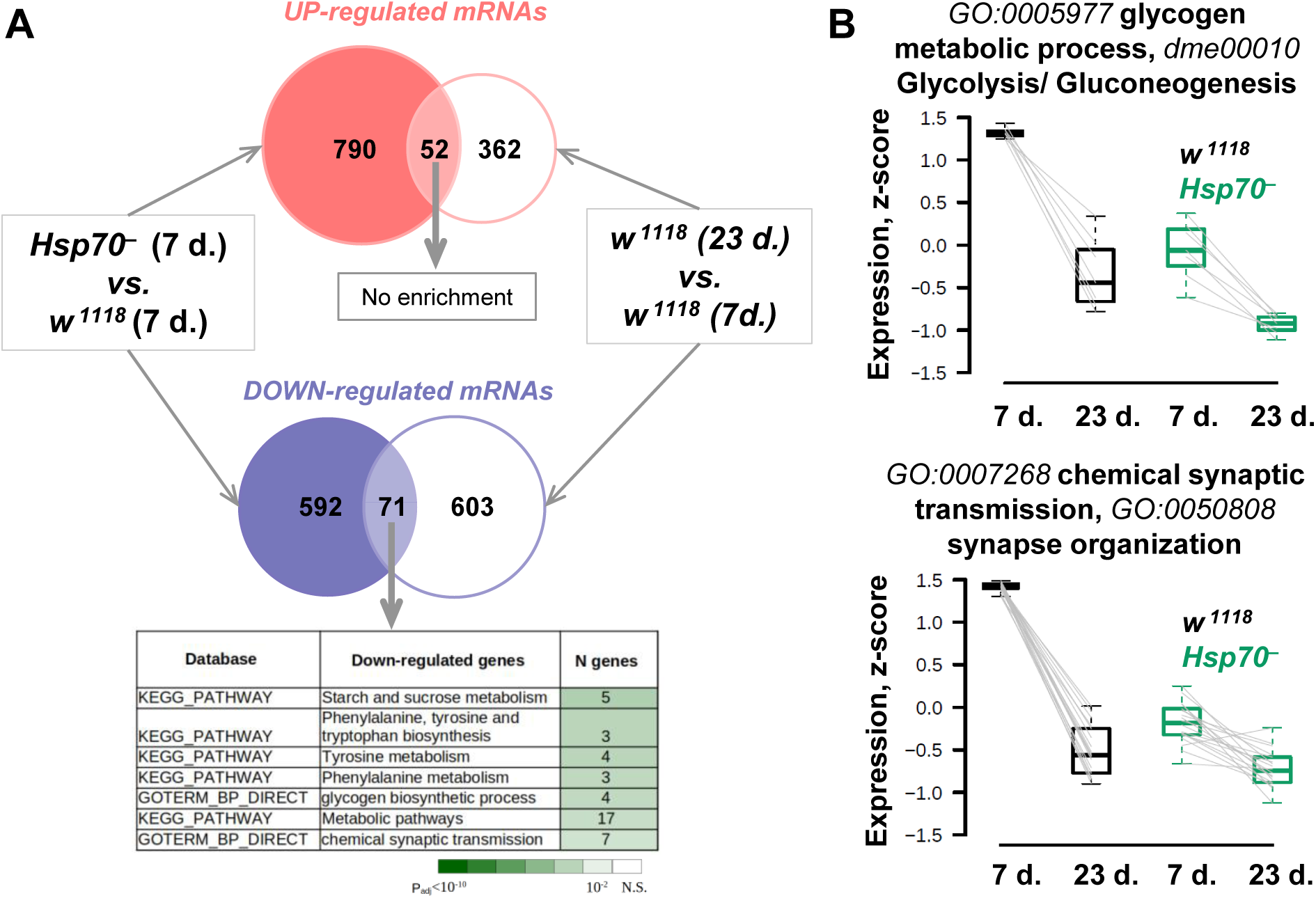
Transcriptomic changes in leg skeletal muscle in 7-day-old *Hsp70^−^* flies are partially associated with age-related changes in *w^1118^* flies. A – Number of up- and down-regulated mRNAs in 7-day-old *Hsp70^−^* flies relative to 7-day-old *w^1118^* and their overlap with age-related changes in *w^1118^* flies (23-day-old *vs.* 7-day-old). n = 4 pools (15 flies per pool) for each strain. An accompanying table shows function of overlapped down-regulated genes. The number of genes in each functional category is presented; the heat map shows the p_adj_-value. B – Expression of genes regulating glycogen metabolism, glycolysis and synapse-related processes in 7- and 23-day-old flies of both strains. Normalized expression (*z*-score) for each differentially expressed gene related to these functional categories (and corresponding IDs) is indicated.

### Knockout of the Hsp70 genes induced widespread changes in leg skeletal muscle transcriptome

Transcriptomic analysis showed that a comparable number of genes changed expression in untrained 7- and 23-day-old *Hsp70^−^* flies relative to the untrained control (600-800 up- and down-regulated mRNAs; Figure 2B). Despite this, significant enrichment of functional categories was found predominantly for 23-day-old flies, which may be associated with age-related progression of specific *Hsp70*-dependent changes. Namely, genes down-regulated in the leg muscles of 23-day-old *Hsp70^−^* flies were associated with glycolysis and the pentose phosphate pathway, amino acid and fat metabolism, as well as with peroxisomal, extracellular, membrane (transporters, membrane-associated metabolic enzymes, G-protein coupled receptors, etc.), and mitochondrial proteins (Supplementary Figure S1). Interestingly, 45 out of 67 down-regulated genes encoding mitochondrial proteins were associated with various mitochondrial enzymes, but there were only a few oxidative metabolism (Krebs cycle and oxidative phosphorylation) enzymes. Up-regulated mRNAs were associated primarily with the extracellular proteins (such as immune humoral factors, chemosensory proteins, chitin- and odorant-binding proteins, and growth factors) and the major regulators of the immune response in *Drosophila* – the Toll and Imd pathway, which activates transcription factor NF-kB (29). Activation of this pathway was indirectly confirmed by the up-regulation of many flies’ antimicrobial peptide genes – potential NF-kB-targets (Supplementary Figure S1). This is in line with data that plasmid-mediated *Hsp70* overexpression in rat skeletal muscle inhibits NF-kB (15).

Altogether, the most striking transcriptomic changes in the 23-day-old *Hsp70^−^* flies are associated with an increase in the expression of genes related to the immune/defense response (including genes encoding extracellular/secretory proteins), as well as with a decrease in the expression of genes encoding regulators of peroxisome and carbohydrate metabolism, and various integral membrane proteins (Supplementary Figure S1).

### Reduced locomotion speed in Hsp70^−^ flies is associated with genotype-specific differences in skeletal muscle gene expression

Interestingly, less than half of the differentially expressed genes in *Hsp70^−^* flies overlapped between days 7 and 23 (Figure 2B), which appears to be due in part to ongoing developmental processes in 7-day-old flies and/or age-related changes in 23-day-old flies. *Hsp70^−^* flies demonstrated a comparable reduction in climbing speed at both time points (Figure 2A); therefore, to identify the functions of genes related to decreased climbing performance in *Hsp70^−^* flies, the overlapped up-regulated (as well as down-regulated) genes were analyzed (Figure 2B). It turned out that the up-regulated genes encoded extracellular proteins (predominantly chitin binding, odorant-binding and chemosensory proteins), regulators of drug metabolism and antioxidant response (cytochromes P450 and glutathione S-transferases), while down-regulated genes encoded regulators of carbohydrate metabolism (in particular, of the pentose phosphate pathway) and various transmembrane proteins (transporters, membrane-associated metabolic enzymes, G-protein coupled receptors, etc.) (Figure 2C).

Then, the enrichment of transcription factor binding sites was examined in the promoter regions of overlapped genes related to decreased locomotion (climbing) performance in *Hsp70^−^*flies. It was revealed that down-regulated genes were associated with transcription factors: Grh (a core component of fibroblast growth factor receptor signaling), Pdm2 (POU-related Homeobox factor) and Trl (regulator of chromatin remodeling), Mad (Bone morphogenetic protein [BMP] signaling; orthologous to human SMAD1/5/9), Atf6 (unfolded protein response signaling), and Adf1; while up-regulated genes with factors: Cf2 (a myogenic factor), Pan (Wnt signaling), Kay (JNK signaling), Hsf (Heat shock factor), and Dve (a repressor involved in chromatin remodeling) (Figure 2D, Supplemental Table S4).

Hsp70 proteins regulate a variety of client proteins (30, 31) and play an important role in regulating age-related changes in proteostasis. Therefore, we examined how changes in the transcriptome caused by knockout of the *Hsp70* genes in 7-day-old flies are comparable with age-related changes in *w^1118^* flies (23 d. vs. 7 d.). Relatively little overlap (52 and 71 up- and down-regulated mRNAs, respectively) were found, but these down-regulated genes enriched terms related to carbohydrate/glycogen and amino acid metabolism, and synaptic transmission – some of the key processes influencing performance of skeletal muscle (Figure 3A and B). Supplementary Figure S2A shows the expression of individual genes related to these terms, as well as genes regulating skeletal muscle contraction. On the other hand, genes regulating DNA repair (*Ku80* and *FANCI*), splicing and translation (*eIF6*, ribosomal protein genes, etc.) were found among the up-regulated overlapped genes; that further characterizes common processes for age-related changes in the control line and in 7 -day-old flies *Hsp70^−^* (Supplementary Figure S2B).

### Disrupted adaptation of Hsp70^−^ flies to chronic exercise training is associated with impaired gene response to exercise in leg skeletal muscles

Adaptation of skeletal muscle to chronic exercise training is associated with transient changes in gene expression after each single exercise bout, as well as with changes in gene expression obtained several days after the last exercise bout (the so-called “baseline” expression) that change muscle proteome and function (13, 32). Chronic climbing training caused small changes (17 mRNAs) in the post-training expression (24 h after the last exercise) in the legs of *w^1118^* flies (Figure 4B), despite an increase in climbing speed induced by chronic training (Figure 4A). Such small changes may be due to the high metabolic rate of flies, i.e., 24-h recovery after the last exercise session could be sufficient to fully restore changes in gene expression induced by chronic training. Knockout of the *Hsp70* genes increased the number of genes related to the post-training expression (Figure 4B). However, in both strains, only a few genes enriched functional categories, which were related to defense and inflammatory response (Supplemental Table S3).

**Figure 4.**
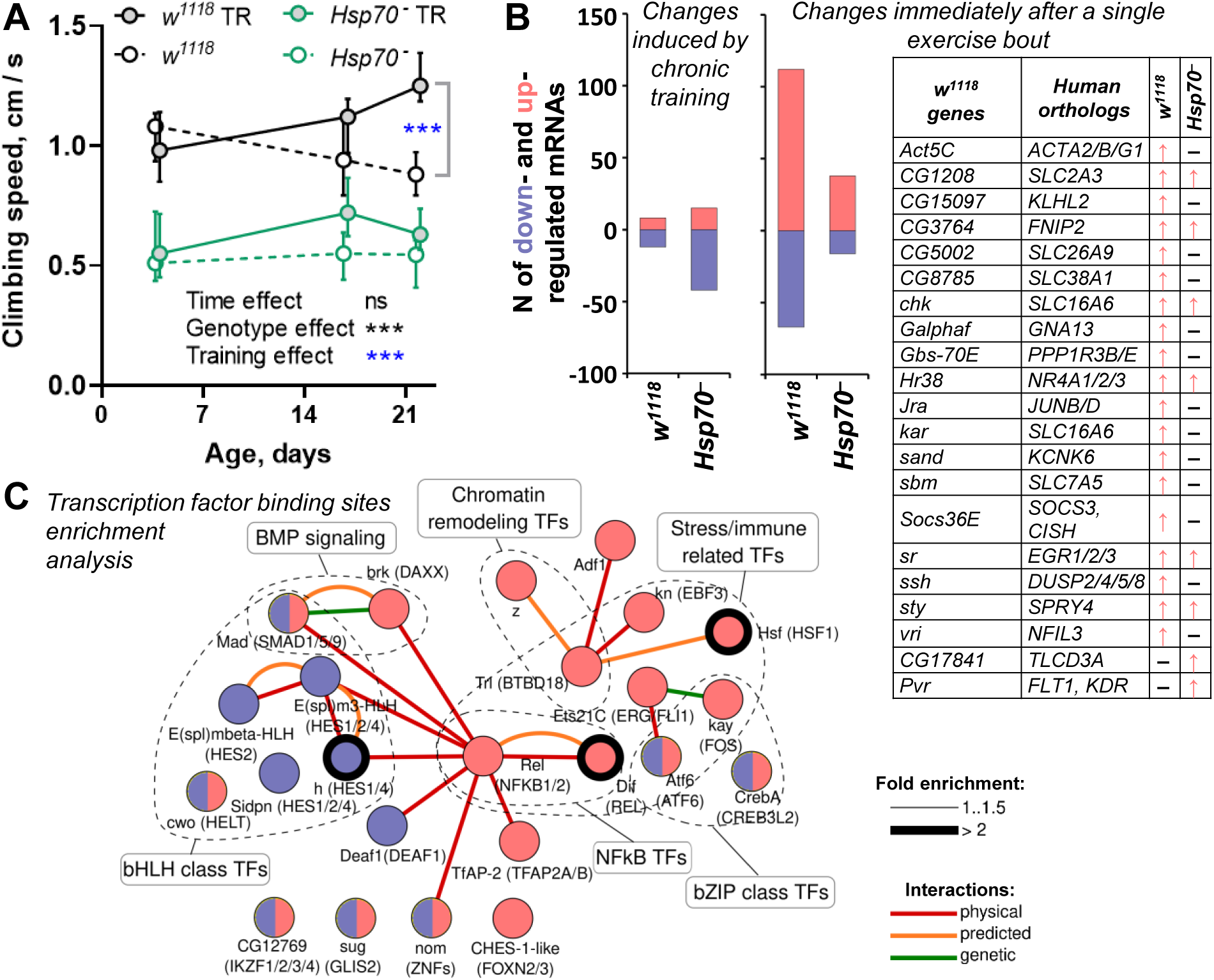
Knockout of six *Hsp70* genes disrupted adaptation of flies to chronic exercise training, which is associated with impaired gene response to a single exercise bout in leg skeletal muscles. A – *Hsp70^−^* flies, in contrast to the control (*w^1118^*), demonstrated no increase in climbing speed after chronic training (4 to 22 days). n = 5–10 pools (20–30 flies per pool) for each strain. TR – training. *** – difference from the control at p <0.001. B – Number of genes changing expression after chronic training (24 h after the last exercise bout) and immediately after a single exercise bout (see Figure 1). Genes responsible to a single exercise bout in *w^1118^*, whose orthologs respond to a single bout of aerobic exercise in human skeletal muscle (according to a meta-analysis (32)), are shown in an accompanying table. Arrows indicate the direction of response in *w^1118^*and human skeletal muscle. n = 4 pools (15 flies per pool) for each strain. C – Transcription factors associated with up- and down-regulated genes (red and blue, respectively) induced by a single exercise bout in the control (*w^1118^*). Lines indicate potential protein-protein and genetic interactions; the border thickness indicates the enrichment of the transcription factor binding site in the promoter (–300 to +100 b.p.).

In contrast, in *w^1118^* flies, a single exercise bout induced expression of 179 genes, which include a number of genes whose orthologs are induced by a single exercise bout in human skeletal muscle (32), namely, transcription factors regulating early gene response (human orthologs: *EGR1*/2/3, *JUNB*, and *NFIL3*), carbohydrate and fat metabolism (*NR4A1/2/3*), STAT-related suppressors of cytokine signaling (*SOCS1/3/6*), transporters (*SLC2A3*, *SLC7A5*, *SLC16A6*, *SLC26A9*, and *SLC38A1*), etc. (Figure 4B). Moreover, a few genes, induced by an exercise, enriched functional categories related to muscle organ and tracheal system development (Supplemental Table S3). Curiously, knockout of the *Hsp70* genes substantially (by 3 times, 54 mRNAs) suppressed genes responsible for a single exercise bout (Figure 4B); additionally, these genes showed no significant enrichment of any functional categories (Supplemental Table S3).

Importantly, in *w^1118^* flies, the transcription factor binding sites enrichment analysis revealed putative transcription factors associated with gene response to a single exercise bout only. Up-regulated mRNAs were associated mainly with factors related to stress and immune response: Rel and Dif (NF-kB transcription factors), Hsf and Atf6 (unfolded protein response signaling), Kay (JNK signaling), Ets21c (an expression marker of various types of cell stress), and Kn (a factor playing a role in muscle development). Meanwhile down-regulated mRNAs were associated with factors of the bHLH family, as well as with some of the factors (Nom, Mad, Creba, Atf6, Cwo, etc.) associated with up-regulated genes (Figure 4C, Supplemental Table S4). Additionally, the analysis of potential protein-protein interactions revealed multiple interactions of transcription factors regulating gene response to a single exercise bout (Figure 4C, Supplemental Table S4), which is in line with findings for human skeletal muscles (32). However, no enrichment of transcription factor binding sites was found for genes induced by both chronic training and a single exercise bout in *Hsp70^−^* flies (Supplemental Table S4).

## Discussion

This study is the first to demonstrate that knockout of *Hsp70* genes not only reduced physical performance, but also disrupted adaptation to chronic physical training, which is associated with changes in skeletal muscle gene expression. The authors analyzed potential mechanisms and revealed that reduced exercise performance in both 7- and 23-day-old *Hsp70^−^* flies was associated with wide-spread change in the transcriptomic profile in the leg skeletal muscle (overlapped differentially expressed genes in the 7- and 23-day-old *Hsp70^−^* flies; Figure 2B) such as: increased expression of genes encoding regulators of drug metabolism [cytochromes P450] and antioxidant response [glutathione S-transferases], as well as decreased expression of genes encoding various transmembrane proteins and enzymes of carbohydrate metabolism [in particular, of the pentose phosphate pathway]. The latter is consistent with a decrease in the content of the pentose phosphate pathway enzymes in the leg muscle of old *Hsp70^−^* flies (23). Moreover, our data showed that transcriptomic changes (and decreased climbing speed) in 7-day-old *Hsp70^−^* flies are partially associated with premature aging, adding to previous findings of age-related changes in these fly strains (23).

The most enriched transcription factors associated with up-regulated genes in *Hsp70^−^* flies were the myogenesis regulator Cf2, an increase in expression of which leads to disruption of the stoichiometry of muscle filaments in adult flies and disruption of the fibril structure (33), and the stress-dependent factor Kay, which negatively affects the function of motor neurons (34). Additionally, the deactivation of transcription factors was predicted: Pdm2, a regulator of motor neuron development (35), Adf1, a regulator of neuronal excitability and synapse maturation, which controls locomotor activity (36), and MAD, a regulator of the development of neuromuscular synapses, the production of neurotransmitters and muscle size (37), which related to the BMP signaling regulating fiber number in the Drosophila adult jump muscle (38). Moreover, in *Hsp70^−^* flies, Hsf activation was predicted, which may be directly caused by the absence of Hsp70 proteins (39).

On the other hand, in the control flies, adaptation to chronic exercise training was associated mainly with gene response to a single exercise bout. It was partially overlapped with that in human skeletal muscle (32) (Figure 4B) indicating that the transient gene response to an exercise is partially conservative in nature. As expected (in line with studies in mammalians (40, 41)), in *w^1118^* flies the most enriched transcription factors associated with up-regulated genes were stress/immune response related factors including Hsf and NF-kB (Dif), as well as the early gene response factors Kay, Atf6, and CrebA (Figure 4C). In contrast, *Hsp70^−^*flies demonstrated significantly impaired gene response to a single exercise bout and no significant enrichment of transcription factors binding sites for genes induced by an exercise that seems to be one of the key mechanisms for disrupting their adaptation to chronic exercise training. Together, the findings of this study complement the evidence that overexpression of the *Hsp70* family gene (*Hspa1a* or *Hspa9*) causes activation of mitochondrial biogenesis in mouse skeletal and cardiac muscle (8, 10) and increases aerobic performance (8).

The change in aerobic performance is closely related to the change in oxidative metabolism and mitochondrial biogenesis in skeletal muscles (4). No decrease was found in the expression of genes encoding the Krebs cycle and oxidative phosphorylation enzymes in the leg muscles of *Hsp70^−^* flies compared to the control, nor an increase in the expression of these genes after chronic training and a single exercise bout in the control (as well as in *Hsp70^−^*) flies. This may be explained primarily by post-transcriptional regulation of the mitochondrial protein expression (reviewed in (4)). Indeed, activation of mitochondrial biogenesis induced by the *Hspa1a* overexpression in mouse skeletal muscle occurs without activation of genes, encoding mitochondrial proteins (8); the same result was obtained when comparing changes in proteomic and transcriptomic profiles in differentiating human progenitor cells (42), mammalian cells responding to misfolding stress (43), and in human skeletal muscle after chronic aerobic training (13, 44). Additionally, the transport of proteins into mitochondria was shown to play an important role in the activation of mitochondrial biogenesis in skeletal muscle (5) and heart (10) induced by chronic contractile activity and overexpression of the *Hspa9*, respectively. It should be noted that chaperones from other families, such as small Hsp (45, 46), Hsp10 and 60 (47, 48), as well as Hsp70 co-chaperone Grpel1 (49), can also regulate mitochondrial biogenesis and metabolic phenotype in muscle cells through the import of mitochondrial proteins.

Our study examined the transcriptomic response in fly legs. It’s important to note, fly legs consisting mainly of muscle bundles and exoskeleton (cuticle) (21), but include many different cells: epithelial, nerve, hemocytes, specialized chemosensory and others (50, 51). Therefore, studying the cell-specific gene expression (for example, using ss/snRNA-seq) seems to be a promising task.In conclusion, the findings of this study clearly indicated that the knockout of the *Hsp70* genes markedly changed the gene expression in leg skeletal muscle and impaired gene response to exercise. Specific mechanisms of *Hsp70*-dependent regulation of gene expression are unclear. The Hsp70 proteins can interact with a variety of client proteins (including ribosomal, nuclear, and mitochondrial proteins) (30, 31) regulating their folding, stability, and transport. The accumulation of unfolded or misfolded proteins may result in the endoplasmic reticulum stress and unfolded protein response (which is in line with activation [predicted] of the Hsf in *Hsp70^−^* flies in the study), thereby inducing expression of many genes. On the other hand, Hsp70s may interact with various signaling proteins and transcription regulators, regulating their stability and activity (52–55) and thereby influencing gene expression, as well as other biological processes. The study of *Hsp70*-dependent mechanisms of gene expression regulation in skeletal muscle, in particular, confirmation of the *Hsp70*-dependent activation/deactivation of transcription factors predicted in this study, seem to be promising tasks for further research. Investigation of these mechanisms may be important for finding ways to prevent muscle functional decline due to disuse, metabolic disorders, and aging.

## Endnotes

RNAseq datasets generated for this study can be found in NCBI Gene Expression Omnibus (GEO) under accession number GSE239395.

## Author Contributions

Conceptualization, P.A.M. and D.V.P.; methodology, P.A.M., I.V.K., and D.V.P.; formal analysis, P.A.M., I.V.K., N.S.K., and D.V.P.; investigation, P.A.M., I.V.K., N.S.K., and D.V.P.; resources, P.A.M. and D.V.P.; data curation, P.A.M. and D.V.P.; writing— original draft preparation, D.V.P.; writing—review and editing, P.A.M., I.V.K., N.S.K., and D.V.P.; project administration, D.V.P. All authors have read and agreed to the published version of the manuscript.

---

**Supplementary Figure S1. Knockout of six *Hsp70* genes induced widespread changes in leg skeletal muscle transcriptome in 7- and 23-day-old flies.**

A – Functional enrichment analysis for up- and down-regulated mRNAs in 7- and 23-day-old *Hsp70^−^* flies compared to the control (*w^1118^*). Significant enrichment of functional categories was found predominantly for 23-day-old flies. The number of genes in each functional category is presented; the heat map shows the p_adj_-value.

B and C – Change in expression of mRNAs encoding integral components of the membrane and proteins of the Toll and Imd pathways and some membrane proteins in the 23-day-old *Hsp70^−^* flies. The number of genes in each protein class is presented (B).

**Supplementary Figure S2. Knockout of six *Hsp70* genes changed age-related gene expression in leg skeletal-muscle.**

A and B – Expression of down- (A) and up-regulated (B) genes found for both: genotype-specific difference in 7-day-old *Hsp70^−^* flies *vs.* 7-day-old *w^1118^*and age-related changes in *w^1118^* flies (23-day-old *vs.* 7-day-old). Frames indicates functional groups: green – «glycogen metabolism/glycolysis», blue – «synaptic transmission», red – «skeletal muscle contraction», brown – «DNA repair», orange – «splicing» and «translation».

**Supplementary Table S1. Transcriptomic changes in leg skeletal muscles of the 7- and 23-day-old *Hsp70^−^* flies relative to *w^1118^* flies (n = 4 pools (15 flies per pool) for each strain).**

**Supplementary Table S2. Putative transcription factors (the position weight matrix approach) associated with transcriptomic changes in the leg skeletal muscles of 7- and 23-day-old *Hsp70^−^*.**

**Supplementary Table S3. Transcriptomic response to chronic climbing training and a single exercise bout in leg skeletal muscles of each strain (*w^1118^* and *Hsp70^−^*) (n = 4 pools (15 flies per pool) for each strain) and function enrichment analysis of differentially expressed mRNAs (p_adj_<0.05, |Fold Change|** ≥**1.25).**

**Supplementary Table S4. Putative transcription factors (the position weight matrix approach) associated with transcriptomic response to chronic climbing training and a single exercise bout in leg skeletal muscles of each strain (*w^1118^* and *Hsp70^−^*).**

