## Supplemental Figure S1 for "Knockout of *Hsp70* genes significantly affects locomotion speed and gene expression in leg skeletal muscles of *Drosophila melanogaster*"

**A**

**DOWN-regulated mRNAs in 7- and 23-day-old Hsp70⁻ flies**

| | GO_DIRECT (BP and CC), KEGG PATHWAY | *Hsp70⁻* vs. $w^{1118}$ 7 day | *Hsp70⁻* vs. $w^{1118}$ 23 day |
|---|---|---|---|
| METABOLIC PATHWAYS | Metabolic pathways | | 139 |
| CARBONHYDRATE METABOLISM | Pentose and glucuronate interconversions | 12 | 20 |
| | Carbon metabolism | | 28 |
| | Pentose phosphate pathway | 7 | 10 |
| | carbohydrate metabolic process | 20 | 26 |
| | Glycolysis / Gluconeogenesis | | 13 |
| | Ascorbate and aldarate metabolism | | 10 |
| AMINOACID METABOLISM | Biosynthesis of amino acids | | 15 |
| | Alanine, aspartate and glutamate metabolism | | 10 |
| | Tyrosine metabolism | | 8 |
| | Phenylalanine metabolism | | 5 |
| | Arginine biosynthesis | | 7 |
| | beta-Alanine metabolism | | 8 |
| | Arginine and proline metabolism | | 8 |
| LIPID METABOLISM | Retinol metabolism | | 10 |
| | Fatty acid degradation | | 9 |
| MITOCHONDRION | mitochondrion | | 67 |
| PEROXISOME | peroxisome (GO_CC) | | 28 |
| | Peroxisome (KEGG) | | 25 |
| DRUG METABOLISM | Metabolism of xenobiotics by cytochrome P450 | | 16 |
| | Drug metabolism - cytochrome P450 | | 14 |
| | Drug metabolism - other enzymes | | 16 |
| EXTRACELLULAR REGION | extracellular space | | 55 |
| | extracellular region | | 53 |
| MEMBRANE | integral component of membrane | 174 | 216 |
| | plasma membrane | 99 | |
| | membrane | 110 | |
| | transmembrane transport | 40 | 44 |
| | integral component of plasma membrane | 43 | |
| SENSORY PERCEPTION | sensory perception of smell | | 16 |

**UP-regulated mRNAs in 7- and 23-day-old Hsp70⁻ flies**

| | GO_DIRECT (BP and CC), KEGG PATHWAY | *Hsp70⁻* vs. $w^{1118}$ 7 day | *Hsp70⁻* vs. $w^{1118}$ 23 day |
|---|---|---|---|
| IMMUNE RESPONSE | defense response to Gram-positive bacterium | | 22 |
| | Toll and Imd signaling pathway | | 22 |
| | antibacterial humoral response | | 14 |
| | response to bacterium | | 17 |
| | defense response | 7 | 15 |
| | defense response to bacterium | | 20 |
| | innate immune response | | 25 |
| | humoral immune response | | 10 |
| | response to wounding | | 10 |
| | defense response to insect | | 6 |
| | defense response to Gram-negative bacterium | | 19 |
| EXTRACELLULAR REGION | extracellular region | 43 | 90 |
| | extracellular space | 30 | 72 |
| SENSORY PERCEPTION | sensory perception of chemical stimulus | 18 | 18 |
| | detection of pheromone | 7 | 7 |
| CUTICLE DEVELOPMENT | chitin-based cuticle development | | 16 |
| DRUG METABOLISM | glutathione metabolic process | | 11 |
| | Metabolism of xenobiotics by cytochrome P450 | 7 | 13 |
| | Drug metabolism - cytochrome P450 | 7 | 13 |
| | Drug metabolism - other enzymes | 8 | 14 |
| | Glutathione metabolism | | 12 |

$P_{adj}<10^{-10}$ … $10^{-2}$ N.S.

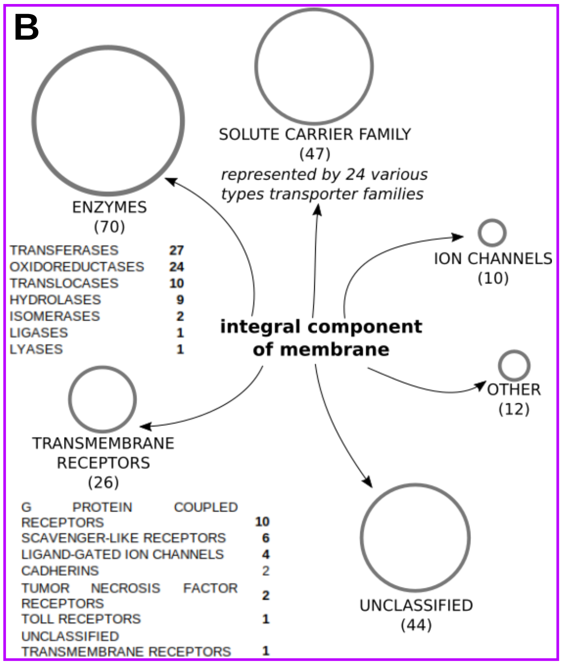


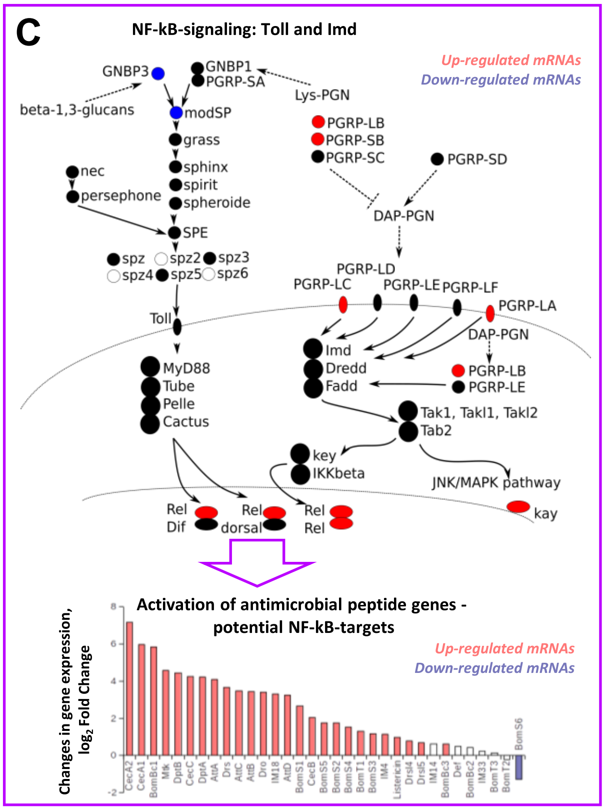


**Supplementary Figure S1. Knockout of six *Hsp70* genes induced widespread changes in leg skeletal muscle transcriptome in 7- and 23-day-old flies.**

A – Functional enrichment analysis for up- and down-regulated mRNAs in 7- and 23-day-old *Hsp70⁻* flies compared to the control ($w^{1118}$). Significant enrichment of functional categories was found predominantly for 23-day-old flies. The number of genes in each functional category is presented; the heat map shows the $p_{adj}$-value.

B and C – Change in expression of mRNAs encoding integral components of the membrane and proteins of the Toll and Imd pathways and some membrane proteins in the 23-day-old *Hsp70⁻* flies. The number of genes in each protein class is presented (B).
