## Supplemental Figure S2 for "Knockout of *Hsp70* genes significantly affects locomotion speed and gene expression in leg skeletal muscles of *Drosophila melanogaster*"

A

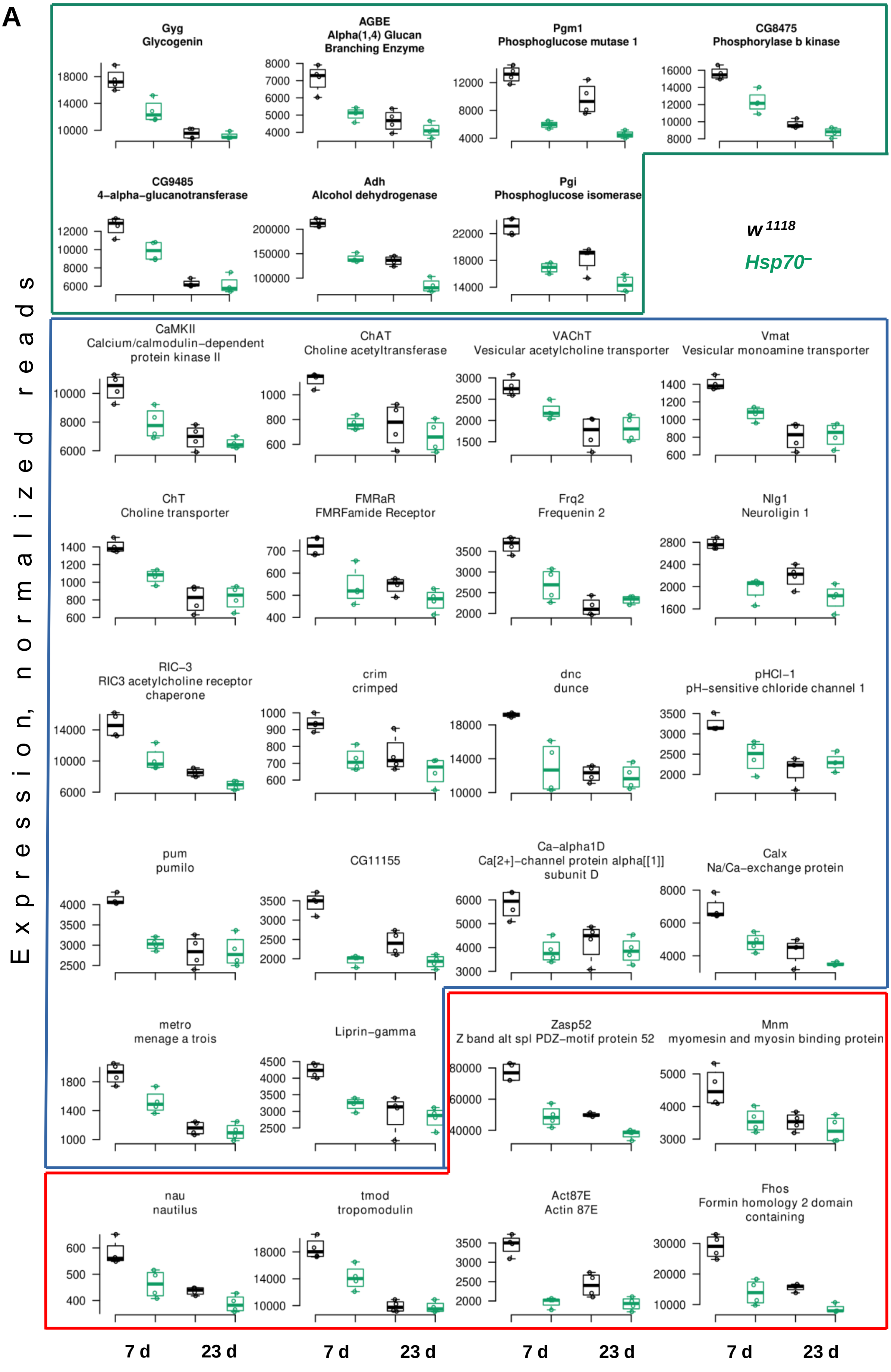

Gyg
Glycogenin
AGBE
Alpha(1,4) Glucan Branching Enzyme
Pgm1
Phosphoglucose mutase 1
CG8475
Phosphorylase b kinase
CG9485
4-alpha-glucanotransferase
Adh
Alcohol dehydrogenase
Pgi
Phosphoglucose isomerase
w 1118
Hsp70–
CaMKII
Calcium/calmodulin-dependent protein kinase II
ChAT
Choline acetyltransferase
VAChT
Vesicular acetylcholine transporter
Vmat
Vesicular monoamine transporter
ChT
Choline transporter
FMRaR
FMRFamide Receptor
Frq2
Frequenin 2
Nlg1
Neuroligin 1
RIC-3
RIC3 acetylcholine receptor chaperone
crim
crimped
dnc
dunce
pHCl-1
pH-sensitive chloride channel 1
pum
pumilo
CG11155
Ca-alpha1D
Ca[2+]-channel protein alpha[[1]] subunit D
Calx
Na/Ca-exchange protein
metro
menage a trois
Liprin-gamma
Zasp52
Z band alt spl PDZ-motif protein 52
Mnm
myomesin and myosin binding protein
nau
nautilus
tmod
tropomodulin
Act87E
Actin 87E
Fhos
Formin homology 2 domain containing
Expression, normalized reads
7 d
23 d

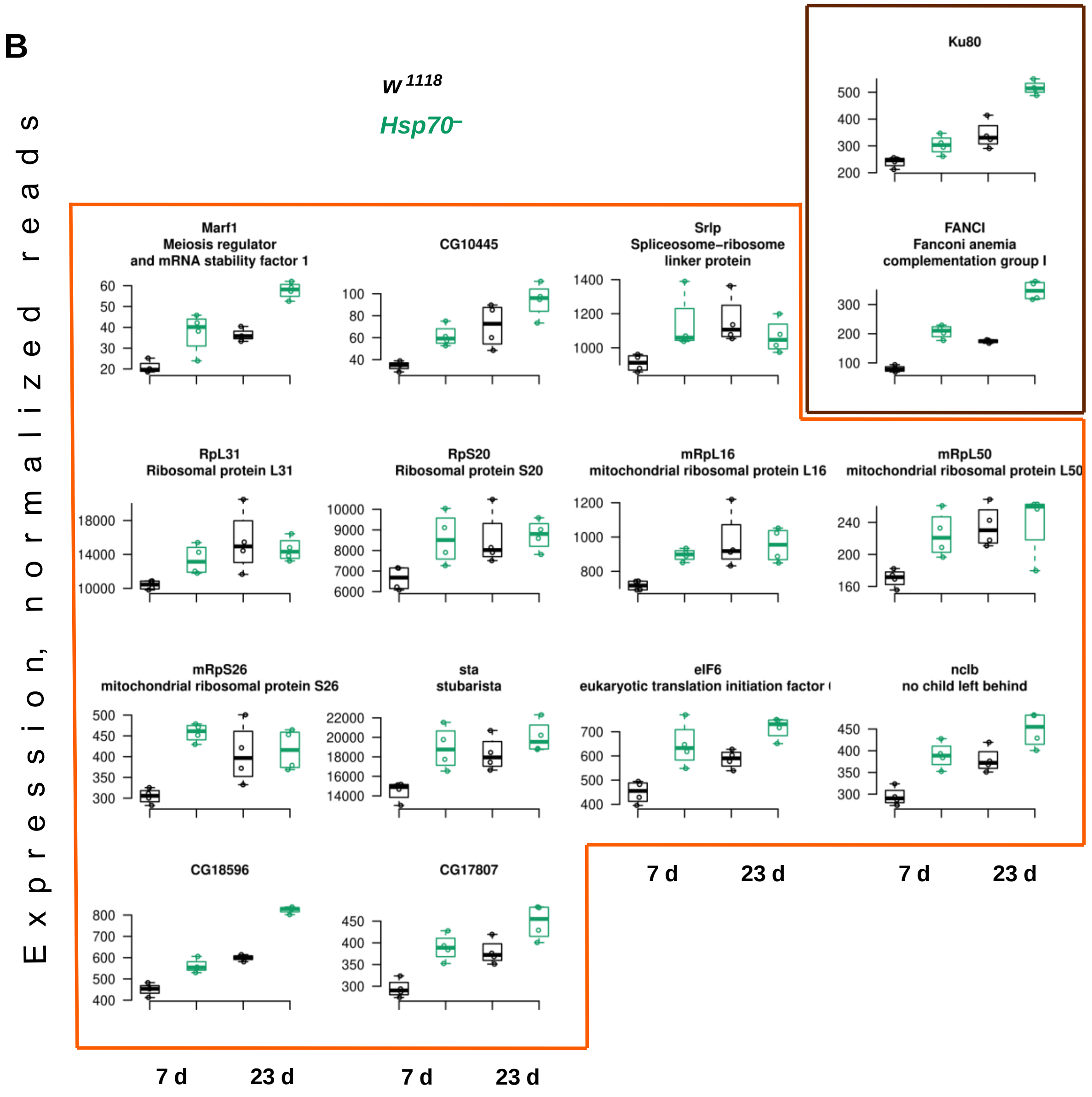


**Supplementary Figure S2. Knockout of six *Hsp70* genes changed age-related gene expression in leg skeletal-muscle.**

A and B – Expression of down- (A) and up-regulated (B) genes found for both: genotype-specific difference in 7-day-old *Hsp70*⁻ flies *vs.* 7-day-old $w^{1118}$ and age-related changes in $w^{1118}$ flies (23-day-old *vs.* 7-day-old). Frames indicates functional groups: green – «glycogen metabolism/glycolysis», blue – «synaptic transmission», red – «skeletal muscle contraction», brown – «DNA repair», orange – «splicing» and «translation».
